# Protein-level prediction of *Klebsiella* phage adsorption identifies conserved receptor-binding motifs

**DOI:** 10.64898/2026.05.21.726843

**Authors:** Francesco Fumagalli, Giacomo Spigler

## Abstract

Bacteriophage therapy offers a potential route to treat antibiotic-resistant *Klebsiella pneumoniae* infections, but its use is limited by the narrow specificity of phage–host interactions. In *Klebsiella*, adsorption is largely determined by receptor-binding proteins (RBPs) that recognize bacterial capsular polysaccharides, yet current machine learning approaches often represent whole phages rather than the individual proteins that mediate recognition. Here, we ask whether adsorption can be predicted at the level of single RBPs and whether the resulting models can identify the molecular features responsible for host specificity.

Using experimentally validated *Klebsiella* phage–host interactions, we extended the PhageHostLearn framework from averaged phage-level representations to individual RBP-level predictions. We found that single-RBP models recover the predictive performance of strain-level models when host capsule identity is explicitly represented. However, models trained only on interaction-level labels did not reliably distinguish motif-bearing RBPs from other viral proteins, indicating that protein-level inputs alone are insufficient for mechanistic interpretability. To resolve this ambiguity, we identified serotype-specific conserved motifs among RBPs from phages infecting the same capsular type. Structural modelling showed that these motifs localize to exposed regions of RBPs and resemble carbohydrate-binding modules. Incorporating motif information into a relabelled training scheme improved prioritization of motif-bearing RBPs while preserving interaction-level predictive power. We further identified a candidate multi-motif RBP from phage S8c that may recognize multiple capsular serotypes.

Together, these results support a modular model of *Klebsiella* phage adsorption in which conserved sub-protein elements drive capsule recognition. More broadly, this work shows how protein-level machine learning combined with biological constraints can move beyond accurate phage–host prediction toward mechanistic identification of host-range determinants.

**Author summary:** Bacteriophages –viruses that infect bacteria– are being explored as alternatives to antibiotics, especially against drug-resistant pathogens such as *Klebsiella pneumoniae*.

The challenge is specificity: each phage attaches to only a narrow range of bacterial strains, recognising them through proteins on its tail that bind the bacterium’s protective sugar capsule. Choosing or engineering the right phage for a given infection therefore requires understanding what these recognition proteins actually do.

We asked whether a machine learning model could move beyond predicting which phages infect a given strain and start identifying which protein on the phage drives that recognition. Prediction alone, we found, is not enough: a model can be accurate without pointing to the responsible protein. To bridge this gap, we searched for short shared sequences among recognition proteins from phages that infect bacteria with the same capsule type, and used these shared patterns to guide the model. This combination correctly prioritised the recognition protein far more often than chance. One phage protein, from phage S8c, carried patterns matching five different capsule types, suggesting a candidate broadly-recognising protein for future experimental study.

## Introduction

Antimicrobial resistance represents one of the most pressing challenges in modern medicine, threatening to undermine decades of progress in treating bacterial infections. Among the pathogens of growing concern is *Klebsiella pneumoniae*, an opportunistic Gram-negative bacterium responsible for a substantial fraction of hospital-acquired infections and increasingly associated with multidrug resistance [1]. A key virulence factor of *Klebsiella* is its polysaccharide capsule, which shields the bacterial surface from environmental threats. These capsules are highly diverse, classified into over one hundred distinct serotypes based on chemical composition [2]. This diversity creates a major obstacle for both antibiotic treatment and alternative therapeutic strategies that depend on precise bacterial targeting.

Bacteriophage therapy has re-emerged as a promising orthogonal approach to combat antibiotic-resistant bacteria [3]. However, its practical deployment is constrained by the high specificity of phage-host interactions. For *Klebsiella*, this specificity is largely determined during adsorption, the initial step of infection, which is mediated by receptor-binding proteins (RBPs) located at the distal end of the viral tail [4, 5]. These proteins recognize specific bacterial surface structures, most notably capsular polysaccharides. Consequently, selecting or engineering therapeutic phages requires identifying RBPs that match the capsule type of a target bacterial strain.

Machine learning (ML) approaches have recently been introduced to predict phage-host interactions directly from genomic data. While several models yield high predictive performance, most operate at a coarse resolution, treating phage and host genomes as monolithic representations [6–8]. PhageHostLearn (PHL) advanced this approach by using embeddings derived explicitly from viral RBPs and bacterial capsule-associated proteins [9]. However, by averaging protein embeddings into a single vector, PHL discards information about individual viral proteins. This limits mechanistic interpretation: such models can predict whether a phage is likely to infect a host, but they do not identify which specific viral component drives adsorption. This is a critical limitation for rational host-range engineering, where the goal is not only to predict infection but also to identify components that can be selected, modified, or transferred [10].

In this work, we address these limitations by shifting the prediction task from whole-phage representations to individual receptor-binding proteins. We first test whether phage adsorption to *Klebsiella* can be predicted from single RBPs without loss of accuracy relative to strain-level models. This allows us to ask whether the predictive signal required for adsorption is encoded within individual viral proteins rather than distributed across whole-phage representations.

We then investigate the molecular basis of this protein-level signal. Specifically, we search for conserved sequence motifs within RBPs from phages targeting the same capsular serotype and map these motifs onto predicted protein structures. These motifs localize to exposed, C-terminal regions, consistent with prior experimental evidence that phage RBPs often have modular architectures and contain carbohydrate-binding modules involved in host recognition [11, 12]. Finally, we test whether incorporating motif information into a relabelled training scheme improves the ability of ML models to prioritize the specific RBPs most likely to mediate adsorption. By combining protein-level ML with domain-specific biological knowledge, this framework links accurate phage-host prediction to mechanistic identification of host-range determinants.

## Results

### Individual receptor-binding proteins recover phage-level adsorption prediction

We first asked whether adsorption of *Klebsiella* phages can be predicted from individual receptor-binding proteins (RBPs), rather than from averaged whole-phage representations. This analysis tests whether the information required for adsorption prediction is concentrated in specific viral proteins or instead depends on a broader phage-level signal.

As a baseline, we replicated the original PhageHostLearn formulation, here referred to as PHL-AVG, in which viral RBP embeddings and bacterial capsule-protein embeddings are averaged into single phage-level and host-level representations. Consistent with the original model, PHL-AVG achieved strong performance at high bacterial genome-similarity thresholds, with performance decreasing as the cross-validation task required generalization to more distantly related bacterial strains (Figure 1).

**Fig 1.**
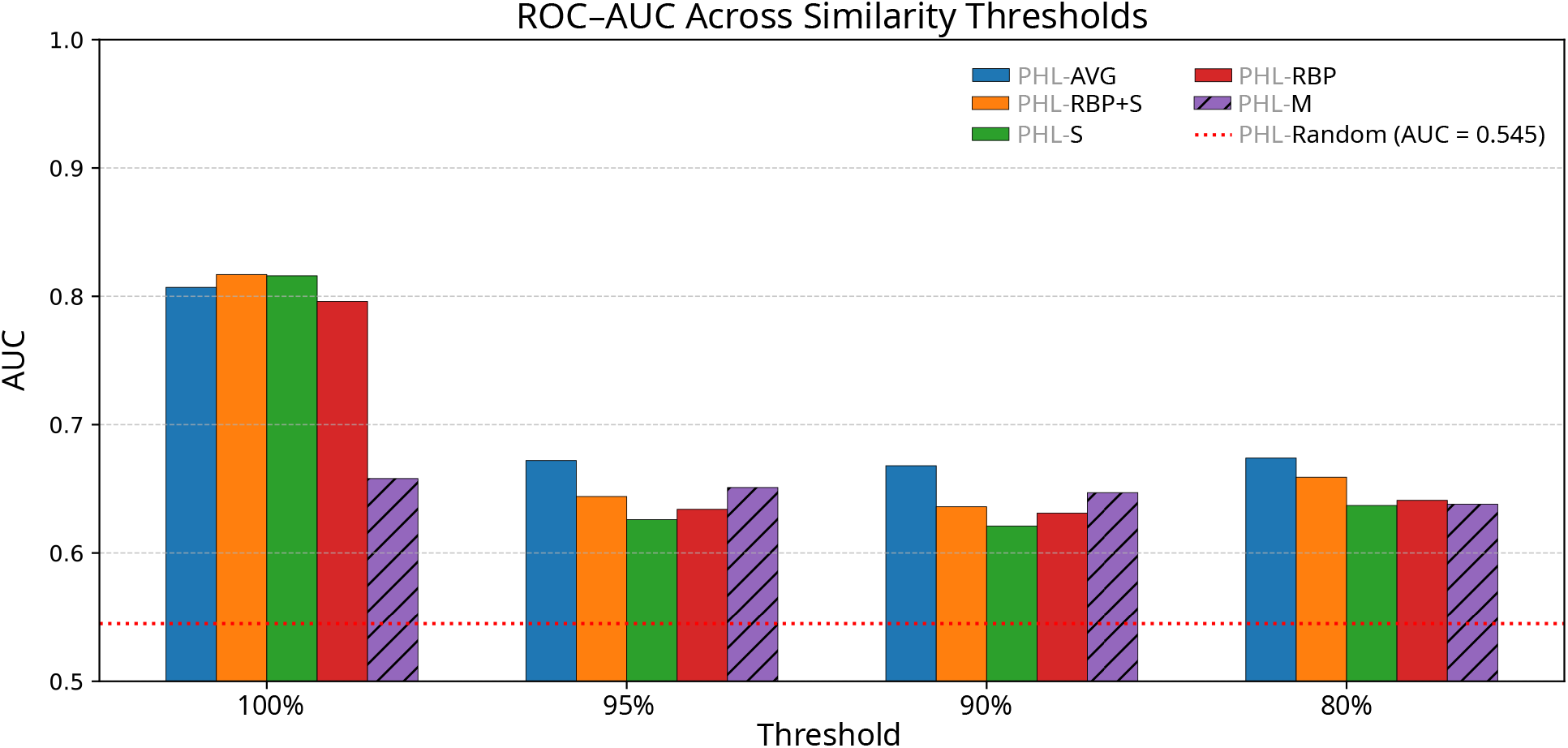
Comparison of phage-host learning strategies across similarity thresholds. Bar plots show model performance (ROC-AUC) evaluated at group-similarity thresholds of 100%, 95%, 90%, and 80%. Models compared include PHL-AVG (averaged phage and host protein embeddings), PHL-RBP (individual viral RBP scores paired with individual host protein embeddings), PHL-S (averaged phage embeddings paired with serotype encoding), and PHL-RBP+S (individual viral RBP scores paired with serotype encoding). PHL-M denotes the motif-informed relabelled model. The dotted red line indicates the highest ROC-AUC obtained across thresholds by a control model trained similarly to PHL-M but using randomly selected proteins rather than motif-bearing RBPs.

We then evaluated a protein-level variant, PHL-RBP, in which individual viral RBPs were scored separately and the maximum RBP-level score was used as the phage-host interaction score. When individual RBPs were paired with individual bacterial capsule-protein embeddings, ROC-AUC remained relatively high, but precision-recall performance decreased substantially. This discrepancy is consistent with the severe class imbalance of the adsorption dataset: naively enumerating individual viral protein-host protein pairings introduces many additional candidate interactions, increasing the false-positive burden even when ranking performance remains acceptable by ROC-AUC.

To reduce this noise while preserving protein-level resolution, we replaced the averaged bacterial capsule-protein embeddings with a one-hot encoding of capsular serotype. This serotype-based representation isolates capsule identity, the host feature most directly linked to *Klebsiella* phage adsorption. When used with averaged viral embeddings, this model, PHL-S, performed similarly to the PHL-AVG baseline and modestly improved precision-recall performance in some similarity-threshold regimes.

Crucially, combining individual viral RBPs with serotype-based host encoding, here referred to as PHL-RBP+S, recovered the performance of the original strain-level model while operating at single-protein resolution. PHL-RBP+S matched PHL-AVG in both ROC-AUC (Figures 1 and 3) and PR-AUC (Figure 2) across the evaluated similarity thresholds. These results indicate that the predictive signal required for *Klebsiella* phage adsorption is retained at the level of individual RBPs, provided that host capsule identity is explicitly represented.

**Fig 2.**
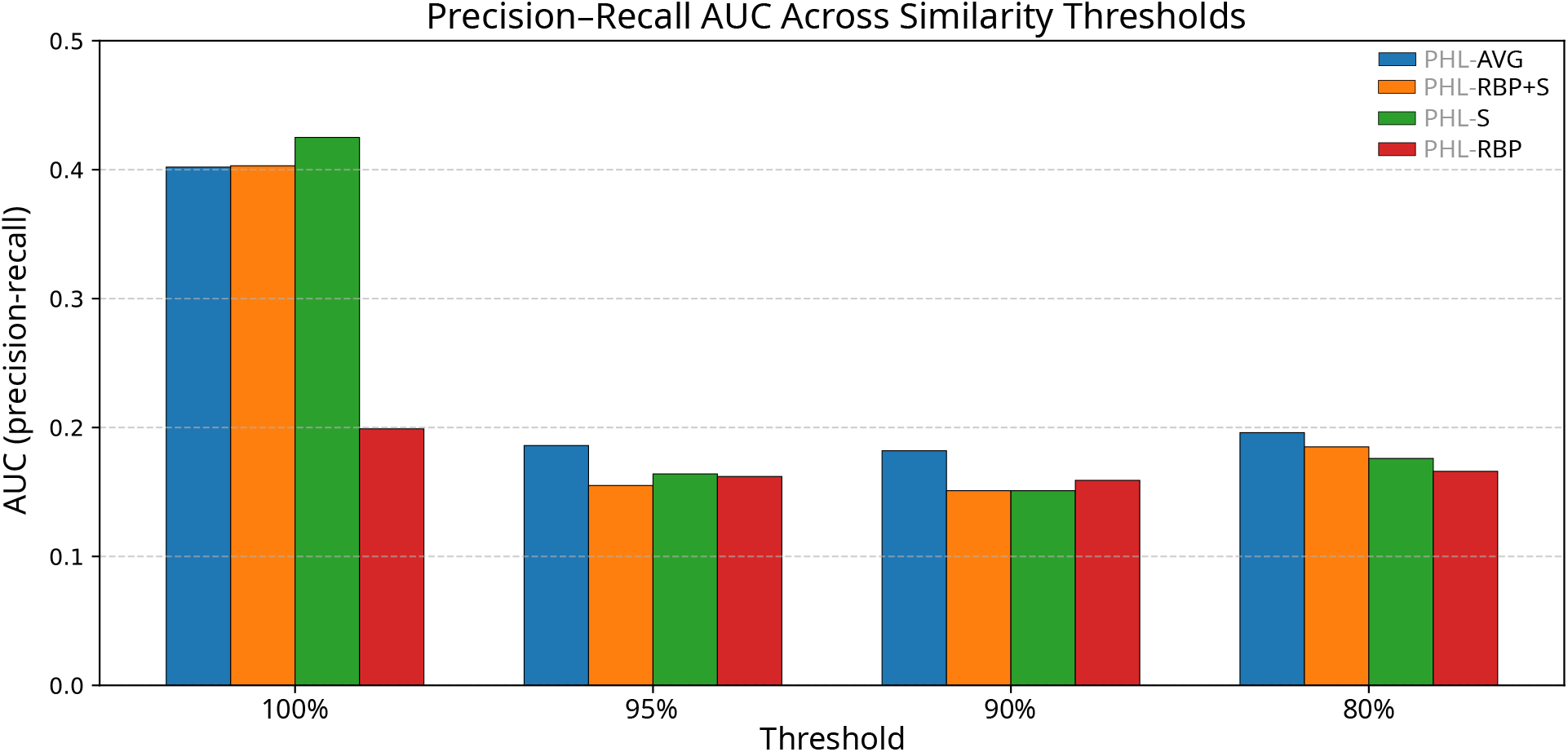
Precision-Recall AUC across group-similarity thresholds for different phage-host interaction models. Bar plots show the area under the precision-recall curve (PR-AUC) for four modelling strategies evaluated at group-similarity thresholds of 100%, 95%, 90%, and 80%.

**Fig 3.**
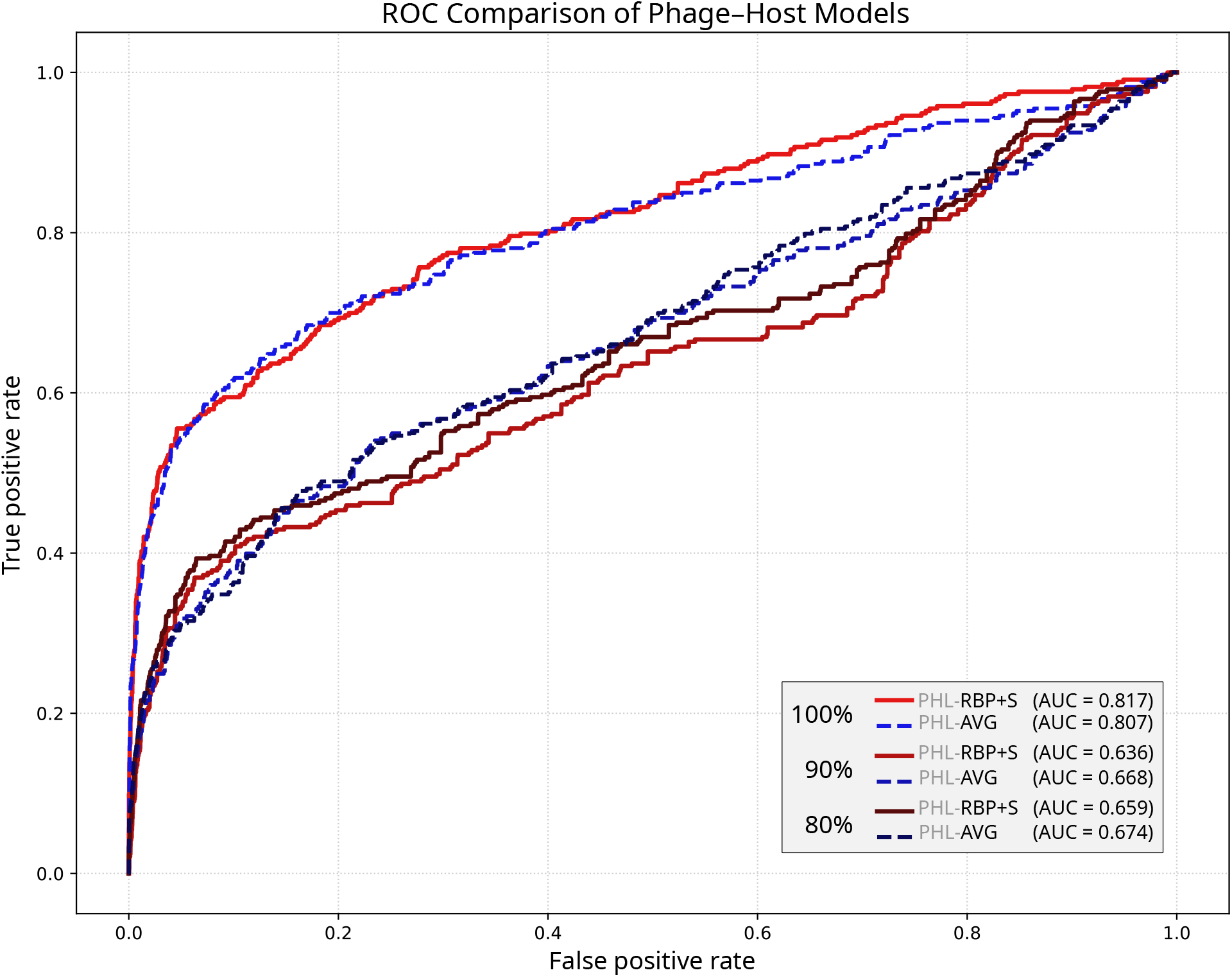
ROC comparison between averaged and individual-protein phage-host models. Receiver operating characteristic (ROC) curves compare the original averaged model (PHL-AVG, dashed blue) with the individual protein-serotype model (PHL-RBP+S, solid red). Curves are shown for group-similarity thresholds of 100%, 90%, and 80%.

### Serotype-specific conserved motifs localize to exposed regions of RBPs

Having established that adsorption can be predicted from individual RBPs, we next investigated whether this protein-level signal could be localized to conserved molecular features. Specifically, we asked whether phages infecting *Klebsiella* strains of the same capsular serotype share conserved RBP elements that could plausibly mediate capsule recognition.

We first tested whether adsorption specificity could be explained by conservation of full-length RBPs. For serotype groups represented by at least six infecting phages, no single full-length RBP was shared across all phages targeting the same serotype. This indicates that common host specificity is not generally explained by identical adsorption proteins, motivating a search for conserved sub-protein features instead.

Guided by the modular architecture of phage RBPs and the known role of carbohydrate-binding modules in host recognition [11, 12], we searched for conserved sequence motifs among RBPs associated with each eligible serotype. For every eligible serotype, motif discovery identified a statistically significant motif shared across the corresponding infecting phages. When mapped onto predicted RBP structures, these motifs consistently localized to C-terminal, surface-exposed regions. In several cases, the highlighted regions protruded from the protein core, consistent with a possible role in capsular polysaccharide recognition (Figure 4; see S3 Appendix for additional serotypes).

**Fig 4.**
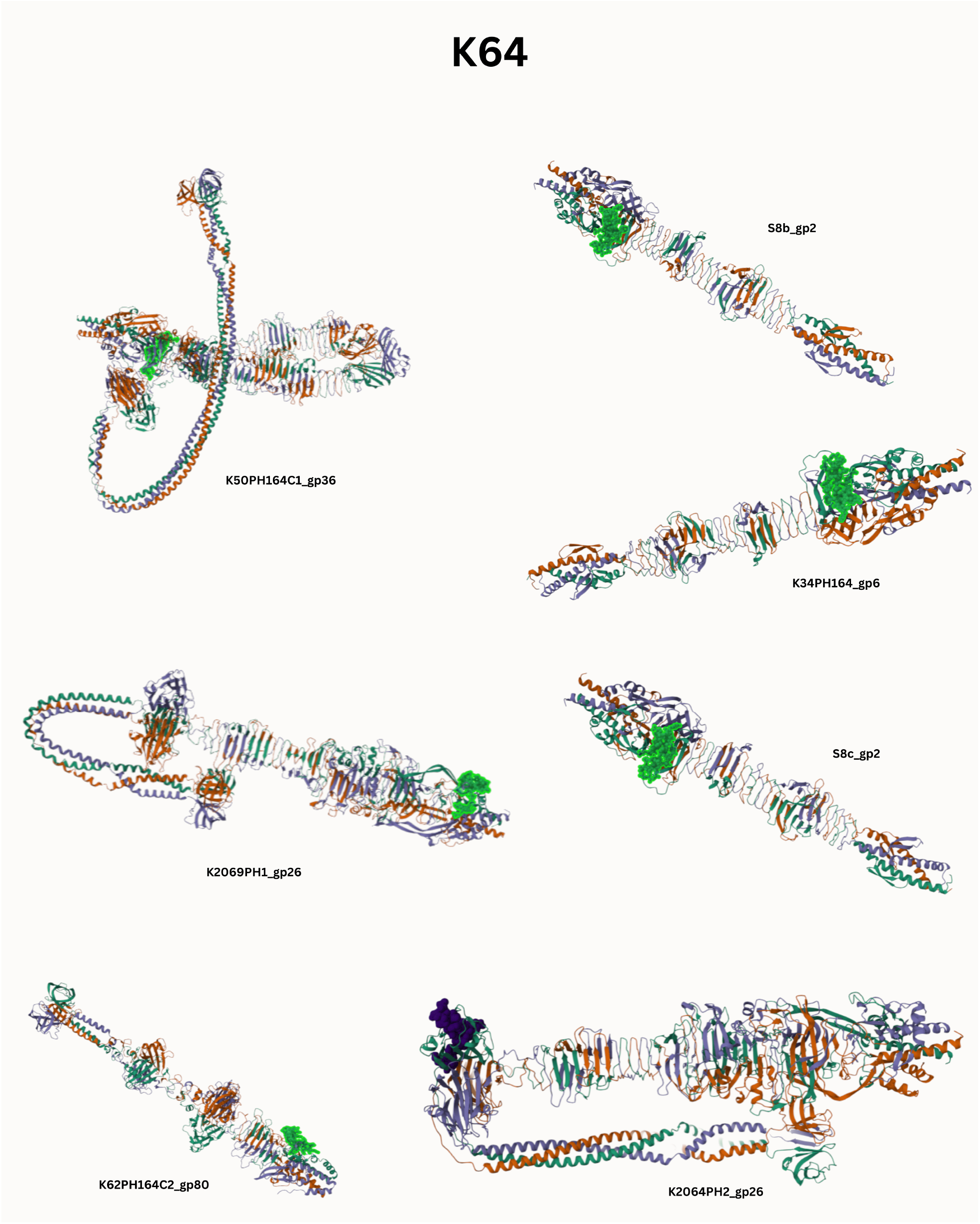
Structural comparison of receptor-binding proteins from K64-infecting phages. Ribbon representations are shown for RBPs from seven distinct phages infecting K64 hosts. A conserved sequence motif, shown in green space-filling representation, is shared across these diverse proteins and consistently maps to exposed structural regions.

These results suggest that, although full-length RBPs vary across phages infecting the same capsular type, shorter conserved motifs may represent shared recognition elements. The convergence of sequence conservation and structural exposure supports the hypothesis that these motifs contribute to adsorption specificity.

### Protein-level prediction alone does not identify the functional RBP

We next asked whether the protein-level model naturally prioritized motif-bearing RBPs over other RBPs encoded by the same phage. This distinction is important because a model may correctly predict that a phage adsorbs to a host without correctly identifying the protein responsible for that interaction.

The standard PHL-RBP+S model did not reliably distinguish motif-bearing RBPs from non-motif-bearing RBPs within the same phage. Predicted scores showed low variance across RBPs associated with the same phage-serotype pair, and motif-bearing RBPs were ranked close to random expectation (S4 Appendix). Thus, although PHL-RBP+S preserved interaction-level predictive performance, its protein-level scores were not sufficient to infer the specific RBP most likely to mediate adsorption.

This result highlights a key interpretability limitation. Moving from averaged phage representations to individual protein inputs increases the resolution of the model input, but does not by itself guarantee mechanistic resolution in the learned predictions. Under interaction-level supervision, predictive signal can remain distributed across correlated proteins from the same phage, even when only one or a subset of those proteins is likely to drive adsorption.

### Motif-informed relabelling improves identification of adsorption-associated RBPs

To test whether the conserved motifs captured functionally relevant information, we trained a motif-informed relabelled model, PHL-M. In this model, motif-bearing RBPs from interacting phage-host pairs were treated as positive protein-level examples, while non-motif-bearing RBPs from those same interacting phages were not used as positives. Interaction-level predictions were still obtained by aggregating RBP-level scores at the phage-host level.

Despite using a more restrictive set of positive protein-level examples, PHL-M retained predictive power on held-out phage-host interactions (Figure 1). More importantly, it substantially improved prioritization of motif-bearing RBPs within multi-RBP phages. At the 100% similarity threshold, motif-bearing RBPs were identified as the most informative protein in 32 of 37 two-RBP cases, 16 of 28 three-RBP cases, and 24 of 42 five-RBP cases, exceeding random expectation in each group (Table 1). These results indicate that the conserved motifs are not merely incidental sequence similarities, but capture information that improves identification of candidate adsorption proteins.

**Table 1.**
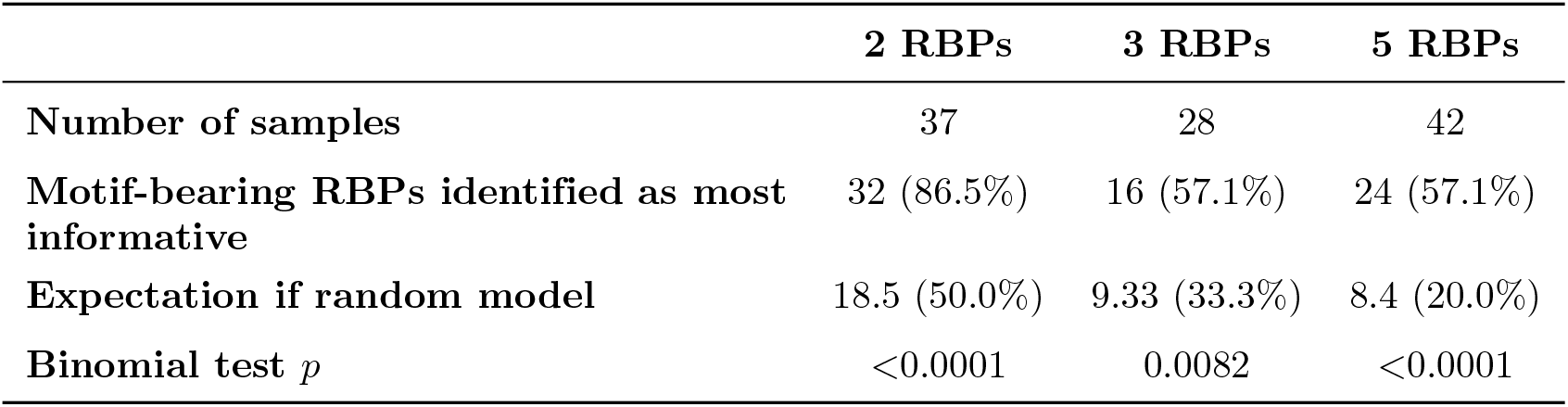
Identification of motif-bearing RBPs by the relabelled model at the 100% genome-similarity threshold.

### A multi-motif RBP suggests a candidate broad-recognition architecture

Finally, we examined whether any individual RBP contained motifs associated with multiple capsular serotypes. One recurrent candidate was gp2 from phage S8c, which was identified as motif-bearing in five distinct serotype groups: K64, K14, K36, K63, and K30 (Figure 5; see S5 Appendix for the full protein and nucleotide sequences). This pattern was not observed for other RBPs in the dataset.

**Fig 5.**
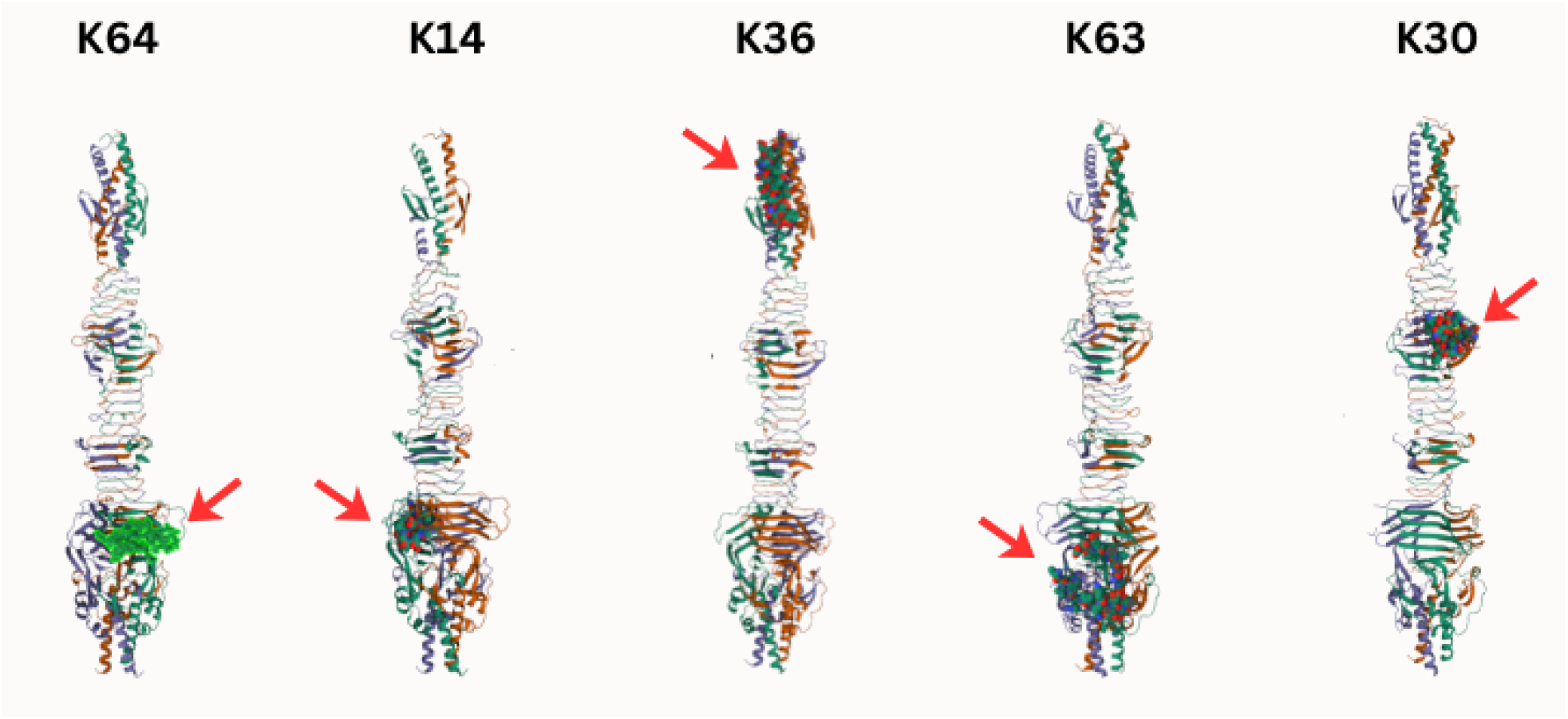
Multi-motif architecture of RBP gp2 from phage S8c. The same protein, gp2, is shown five times. In each instance, a different structural region is highlighted in space-filling representation, corresponding to a conserved motif shared with phages infecting a specific serotype: K64, K14, K36, K63, or K30. This illustrates a single protein containing multiple distinct candidate recognition modules. Red arrows mark the location of the highlighted motif in each case.

Structural mapping suggests that the motifs associated with gp2 occupy distinct regions of the same protein, consistent with a possible multi-domain or multi-module recognition architecture. This makes gp2 a candidate for future experimental validation as a broad-recognition RBP. However, because the present study is computational, its potential role in expanded host range should be interpreted as a hypothesis rather than a demonstrated functional property.

## Discussion

In this study, we investigated the molecular determinants of host specificity in *Klebsiella* phages by combining protein-level machine learning with sequence- and structure-based motif analysis. We show that adsorption can be predicted from individual receptor-binding proteins (RBPs) without loss of performance relative to averaged phage-level representations, provided that host capsule identity is explicitly represented. This finding supports a modular view of *Klebsiella* phage adsorption, in which much of the predictive signal for host recognition is concentrated within individual RBPs rather than distributed across whole-phage representations.

A central implication of this result is that phage-host prediction can be made more biologically interpretable without necessarily sacrificing performance. Previous genomic machine learning approaches have achieved strong predictive accuracy, but often rely on whole-genome or averaged representations that obscure the specific viral components responsible for adsorption [6–8]. PhageHostLearn improved biological specificity by incorporating viral RBP and bacterial capsule-protein embeddings [9], but its averaging strategy still collapsed multiple viral proteins into a single phage-level representation. By scoring individual RBPs directly, our approach preserves protein-level information while retaining interaction-level predictive accuracy.

However, our results also show that protein-level inputs alone are not sufficient for mechanistic interpretation. Although the PHL-RBP+S model operated on individual RBPs, it did not reliably distinguish motif-bearing RBPs from other RBPs encoded by the same phage. This suggests that, under interaction-level supervision, the model can assign similar predictive scores to proteins that are correlated at the genome level, even if only one or a subset of those proteins is likely to mediate adsorption. In practical terms, this means that increasing the granularity of the input representation does not automatically produce a mechanistic explanation. For phage engineering applications, this distinction is critical: predicting that a phage adsorbs to a host is not equivalent to identifying the protein element that determines host range.

To address this limitation, we incorporated domain-specific biological information through conserved motif discovery. Full-length RBPs were not generally conserved across phages infecting the same capsular serotype, but shorter serotype-associated motifs were consistently identified. When mapped onto predicted structures, these motifs localized to exposed, often C-terminal regions of RBPs. This pattern is consistent with previous experimental and structural studies showing that phage RBPs often have modular architectures and contain carbohydrate-binding regions involved in host recognition [11, 12]. The convergence of sequence conservation, structural exposure, and improved model prioritization supports the hypothesis that these motifs represent candidate determinants of capsule recognition.

The motif-informed relabelling experiment further demonstrates the value of incorporating biological constraints into machine learning models. By treating motif-bearing RBPs as the relevant positive protein-level examples, the relabelled model better prioritized candidate adsorption proteins while retaining predictive power at the phage-host level. This result suggests that conserved motifs capture functionally relevant information that is not fully recovered from interaction labels alone. More broadly, it illustrates a useful strategy for interpretable biological prediction: weak interaction-level labels can be complemented with domain-informed intermediate labels to guide models toward more mechanistically meaningful features.

From an applied perspective, identifying conserved RBP motifs may help guide the selection or engineering of phages with defined host ranges. In *Klebsiella*, where capsule type is a major determinant of adsorption, motif-level information could help identify candidate RBPs for experimental screening, host-range retargeting, or modular engineering. The recurrent identification of gp2 from phage S8c as motif-bearing across multiple serotype groups is particularly notable. Structural mapping suggests that different motifs occupy distinct regions of this protein, raising the possibility of a multi-module recognition architecture. However, this interpretation remains computational. Direct experimental validation will be required to determine whether gp2 can mediate adsorption to multiple capsular types and whether its motifs can be transferred or modified to alter host range.

Several limitations should be acknowledged. First, our analyses are based on a single curated dataset of *Klebsiella* phage-host interactions. Although this dataset is well suited for benchmarking against PhageHostLearn, the generality of the approach to other bacterial species, capsule systems, or receptor classes remains to be established. Second, the serotype-based host representation assumes that capsule identity is known and accurately assigned. This is reasonable for many clinically relevant *Klebsiella* isolates, but it may limit application to strains with poorly characterized or unusual capsule loci. Third, the structural analyses rely on predicted protein structures rather than experimentally resolved RBP structures. AlphaFold-based mapping is useful for hypothesis generation, but surface exposure and motif placement should ultimately be confirmed experimentally. Finally, the motif-informed labels used here are computationally inferred, meaning that the relabelled model prioritizes candidate functional RBPs rather than experimentally proven adsorption determinants.

Future work should therefore focus on experimental validation of the predicted motifs and candidate multi-motif RBPs. Targeted mutagenesis, motif deletion, domain swapping, and adsorption assays could directly test whether the identified motifs are necessary or sufficient for capsule recognition. Expanding the approach to additional phage-host systems would also clarify whether motif-informed protein-level prediction is specific to *Klebsiella* capsule recognition or represents a more general framework for identifying host-range determinants.

Overall, this study shows that accurate phage-host prediction can be linked to mechanistic hypotheses by moving from whole-phage representations to individual proteins and then from individual proteins to conserved sub-protein motifs. For *Klebsiella* phages, these results support a model in which adsorption specificity is governed by modular RBP elements that recognize capsular polysaccharides. By combining protein-level machine learning with motif discovery and structural interpretation, this framework provides a route toward more interpretable prediction of phage host range and a starting point for rational phage engineering.

## Materials and Methods

### Dataset and evaluation protocol

We based our analyses on an existing dataset of experimentally validated phage-host interactions for *Klebsiella* spp. The dataset comprises 105 bacteriophage genomes, 200 bacterial host genomes corresponding to distinct strains, and a matrix of 10,006 observed phage-host pairs, of which 3.33% represent confirmed interactions [13, 14]. Positive interactions were identified *in vitro* by the presence of plaques at a 1:10 phage dilution, resulting in a highly imbalanced classification problem.

To evaluate model generalization to unseen strains while preventing information leakage from genomic similarity, we adopted the Leave-One-Group-Out (LOGO) cross-validation strategy introduced in PhageHostLearn [9]. Bacterial genomes were grouped by pairwise genomic similarity, with thresholds ranging from 100% (identical genomes at K-loci) down to 75% (see S1 Appendix). For each threshold, models were trained on all but one bacterial group and evaluated on the excluded group, ensuring each interaction appeared exactly once in a validation set. Performance is reported as the aggregate across all held-out groups.

Given the severe class imbalance and the high downstream cost of false positives in phage therapy, we primarily assessed performance using precision-recall (PR) curves and the area under the curve (PR-AUC) [15]. Receiver operating characteristic (ROC) curves are reported for comparison with prior work. Detailed curves for all model variants and similarity thresholds are provided in S2 Appendix.

### Protein-level extension of PhageHostLearn

Our baseline was the PhageHostLearn (PHL) pipeline [9], which predicts adsorption using embeddings of viral receptor-binding proteins (RBPs) and bacterial capsule-related proteins. In the original PHL formulation, RBPs are identified from phage genomes using Phanotate [16], while capsule-associated proteins are extracted from bacterial genomes using Kaptive [17]. Then, 1280-dimensional embeddings (generated via ESM-2 [18]) of the respective proteins are averaged to create monolithic representations for each phage and host.

To retain protein-level resolution, we introduced two modifications. First, rather than averaging viral RBP embeddings, we treated each RBP as a distinct entity during training and inference. Depending on the model variant, these viral embeddings were paired either with individual bacterial capsule-protein embeddings, or with a reduced host representation (serotype encoding). This modification allowed the models to operate at the level of individual viral proteins while preserving the overall prediction task of phage-host adsorption.

For phages expressing multiple RBPs, inference produces one score per viral protein-capsule protein pairing. To derive a single interaction score for the phage-host pair, we retained the maximum score across all combinations with RBPs expressed by that phage and that capsule. This aggregation strategy reflects the biological assumption that successful adsorption can be driven by a single key protein, consistent with modular phage tail architectures.

All models were trained using XGBoost with the hyperparameters reported in the original PHL study to ensure direct comparability (reported in S1 Appendix). No additional hyperparameter tuning was performed unless explicitly stated. For every training instance, the viral representation (either the averaged phage embedding for the baseline or a single RBP embedding for our extension) was concatenated with the corresponding host representation to form the input vector.

### Capsular serotype encoding

Complementing the protein-level host representations, we explored a reduced encoding based solely on capsular serotype. Using Kaptive [17], each bacterial genome was assigned a K-locus corresponding to its capsular chemical composition. This information was encoded as a one-hot vector and used in place of the averaged capsule-protein embeddings.

This serotype-based encoding substantially reduces input dimensionality and isolates the specific contribution of capsule identity to adsorption. While this limits applicability to hosts with characterized capsules, the vast majority of clinically relevant *Klebsiella* isolates belong to known serotypes [19], rendering this limitation minor in a therapeutic context.

### Motif discovery and structural mapping

To investigate the molecular basis of the predictive signal, we searched for conserved sequence motifs among RBPs from phages infecting the same capsular serotype. For each serotype associated with at least six infecting phages, we collected all RBPs expressed by those phages and compared them. Motif discovery was attempted for all serotypes with at least three infecting phages; structural mapping and model training were restricted to the eight serotypes with at least six infecting phages (K11, K13, K14, K29, K30, K36, K63, K64), for which sufficient sequence diversity was present.

Motif identification was performed using MEME (Multiple EM for Motif Elicitation) [20]. For each serotype, we retained the most statistically significant motif (length 21-99 amino acids). To assess structural plausibility, we generated three-dimensional structures of trimers from the corresponding RBPs using AlphaFold3 [21] and mapped the conserved motifs onto these structures using Mol* Viewer [22]. This allowed us to inspect motif localization, surface exposure, and spatial consistency.

### Protein-level scoring and relabelled training

We quantified the importance of motif-bearing RBPs using two approaches. First, we used the protein-serotype model to score individual RBPs under LOGO cross-validation. Since each RBP was scored by a model not trained on its associated group, we could unbiasedly rank the proteins within each phage-host pair to see if motif-bearing RBPs were prioritized.

Second, we implemented a “relabelled” training scheme to explicitly test the motif hypothesis. In this regime, we modified the training labels: only RBPs containing a serotype-specific conserved motif were assigned positive labels, whereas non-motif-bearing RBPs from interacting phages were ignored. Negative samples were drawn from non-interacting phage-host pairs. Models trained under this scheme still produced interaction-level predictions via maximum-score aggregation and were evaluated against the original experimentally validated interaction labels. This experiment tests whether strictly supervising the model to look for motifs improves its ability to identify valid phage-host pairs.

### Statistical comparison with random expectation

For phages encoding multiple RBPs, we compared the number of cases in which the model ranked a motif-bearing RBP highest against the number expected under random selection. Random expectation was calculated as 1*/n*, where *n* is the number of RBPs encoded by the phage, aggregated across all phage-host cases in each RBP-count group.

Statistical significance was assessed using binomial tests comparing the observed number of motif-bearing RBPs ranked highest with the random expectation for that group. Results were summarized separately for phages encoding different numbers of RBPs.

## Data and code availability

The dataset used in this study is available through the Zenodo repository associated with Boeckaerts et al. [9]. The code required to reproduce the analyses, figures, model hyperparameters, motif discovery outputs, and trained protein-level models is available at: https://github.com/FumaNet/TheKeyMotif.git

## Acknowledgments

The authors thank Dimitri Boeckaerts and colleagues for making their dataset and code publicly available, and for helpful discussions about their work.

## Supporting information

**S1 Appendix. Genome similarity thresholds and XGBoost hyperparameters**. This appendix reports the bacterial genome similarity groupings used for Leave-One-Group-Out cross-validation and the XGBoost hyperparameters used across classifiers. These details support reproducibility of the evaluation protocol and ensure comparability between the original PhageHostLearn baseline and the protein-level model variants.

**Table 2.**
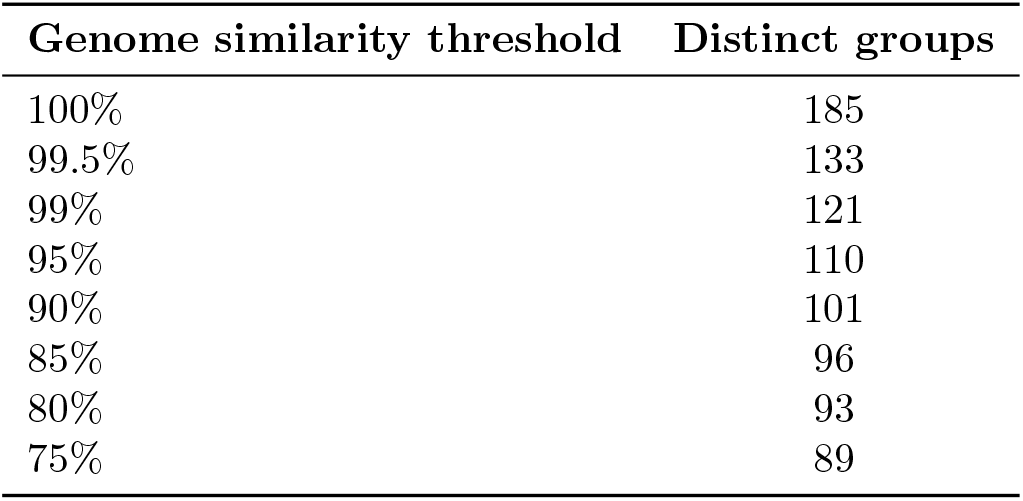
Number of bacterial genome similarity groups formed at different genome similarity thresholds. The dataset contains 200 bacterial genomes.

| Genome similarity threshold | Distinct groups |
| --- | --- |
| 100% | 185 |
| 99.5% | 133 |
| 99% | 121 |
| 95% | 110 |
| 90% | 101 |
| 85% | 96 |
| 80% | 93 |
| 75% | 89 |

**Table 3.** XGBoost hyperparameters used for all classifiers in this study. Values correspond to the optimal configuration reported by Boeckaerts et al. for PhageHostLearn and were retained across XGBoost-based classifiers for consistency and comparability.

| Hyperparameter | Value |
| --- | --- |
| Maximum tree depth | 7 |
| Learning rate | 0.3 |
| Number of estimators | 250 |

**S2 Appendix. ROC and precision-recall curves for PhageHostLearn model variants**. This appendix provides the complete ROC and precision-recall curves for each model variant across the genome similarity thresholds used in Leave-One-Group-Out cross-validation. These plots supplement the main-text AUC summaries by showing model behavior across the full decision-threshold range.

**Fig 6.**
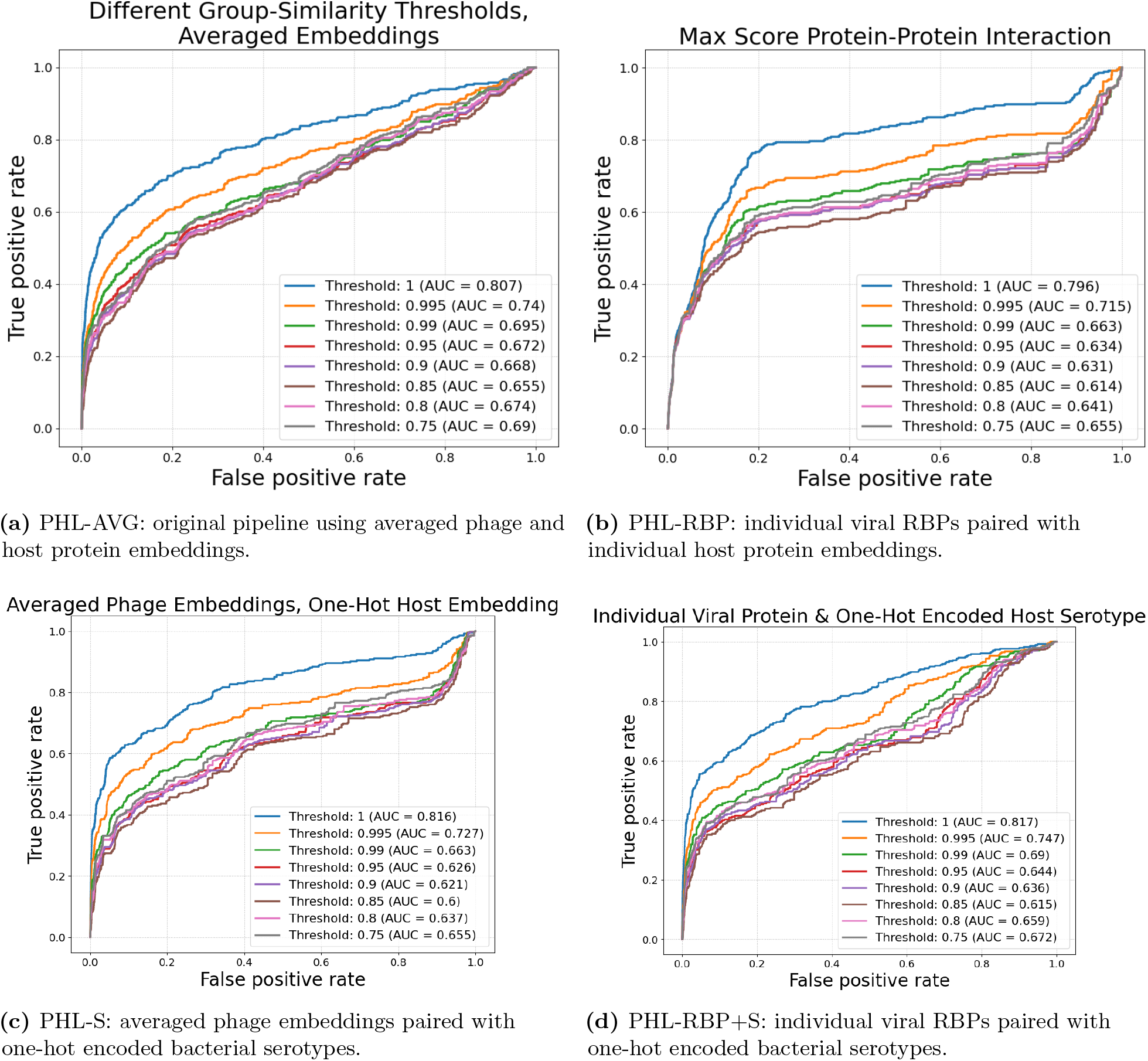
ROC curves for the baseline PhageHostLearn model and protein-level extensions across LOGO cross-validation similarity thresholds.

**Fig 7.**
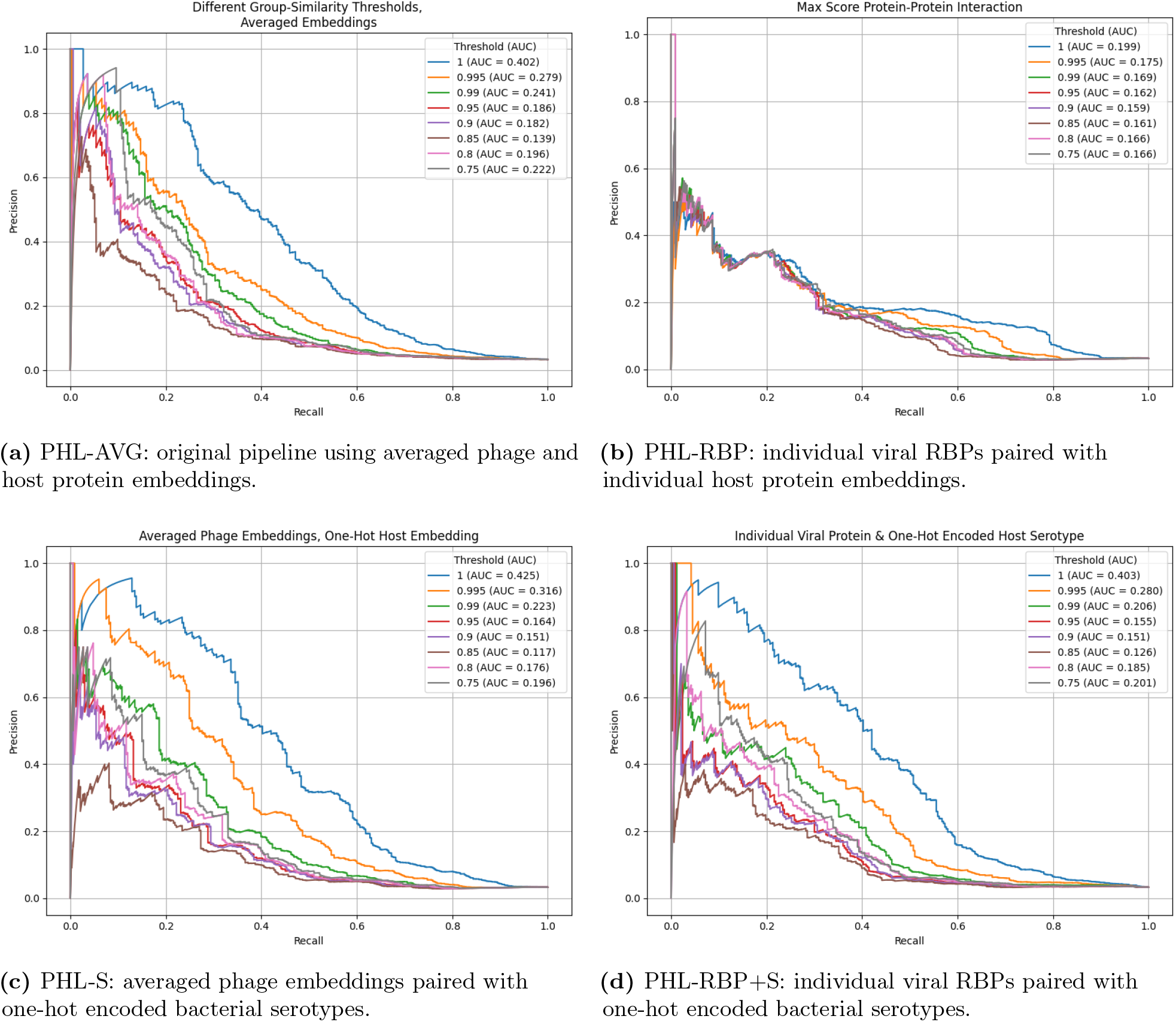
Precision-recall curves for the baseline PhageHostLearn model and protein-level extensions across LOGO cross-validation similarity thresholds.

**Fig 8.**
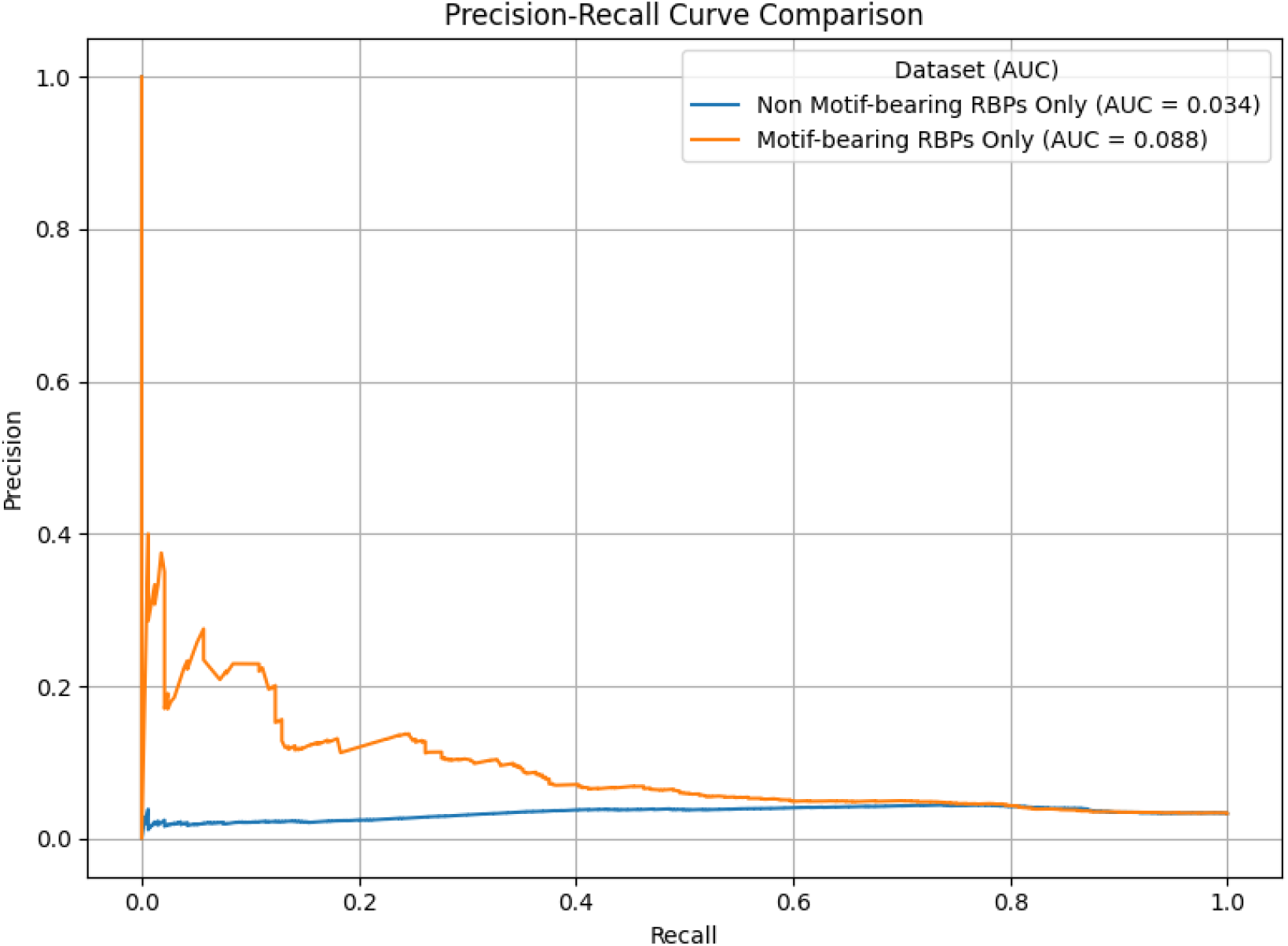
Precision-recall AUC comparison between models trained on motif-bearing RBPs and control models trained on non-motif-bearing RBPs at the 100% LOGO threshold.

**S3 Appendix. Structural mapping of serotype-associated conserved motifs**. This appendix shows AlphaFold-predicted structures for RBPs containing conserved motifs associated with specific *Klebsiella* capsular serotypes. Highlighted regions indicate motif positions within each predicted structure, allowing comparison of motif localization and surface exposure across serotype groups.

**Fig 9.**
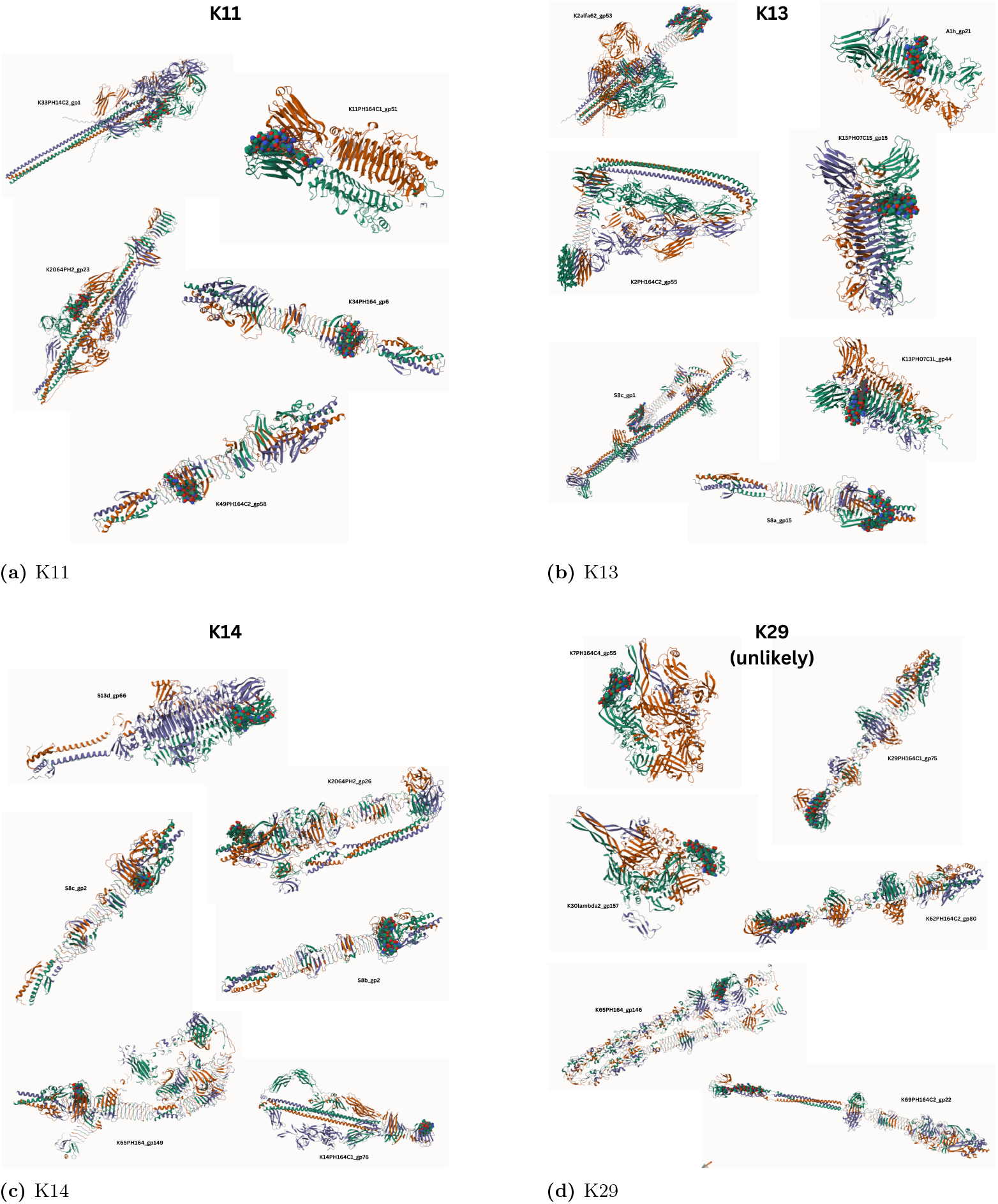
Predicted structures of RBPs containing conserved serotype-associated motifs for K11, K13, K14, and K29. Motifs are shown in molecular representation, replacing the ribbon representation used for the rest of each protein.

**Fig 10.**
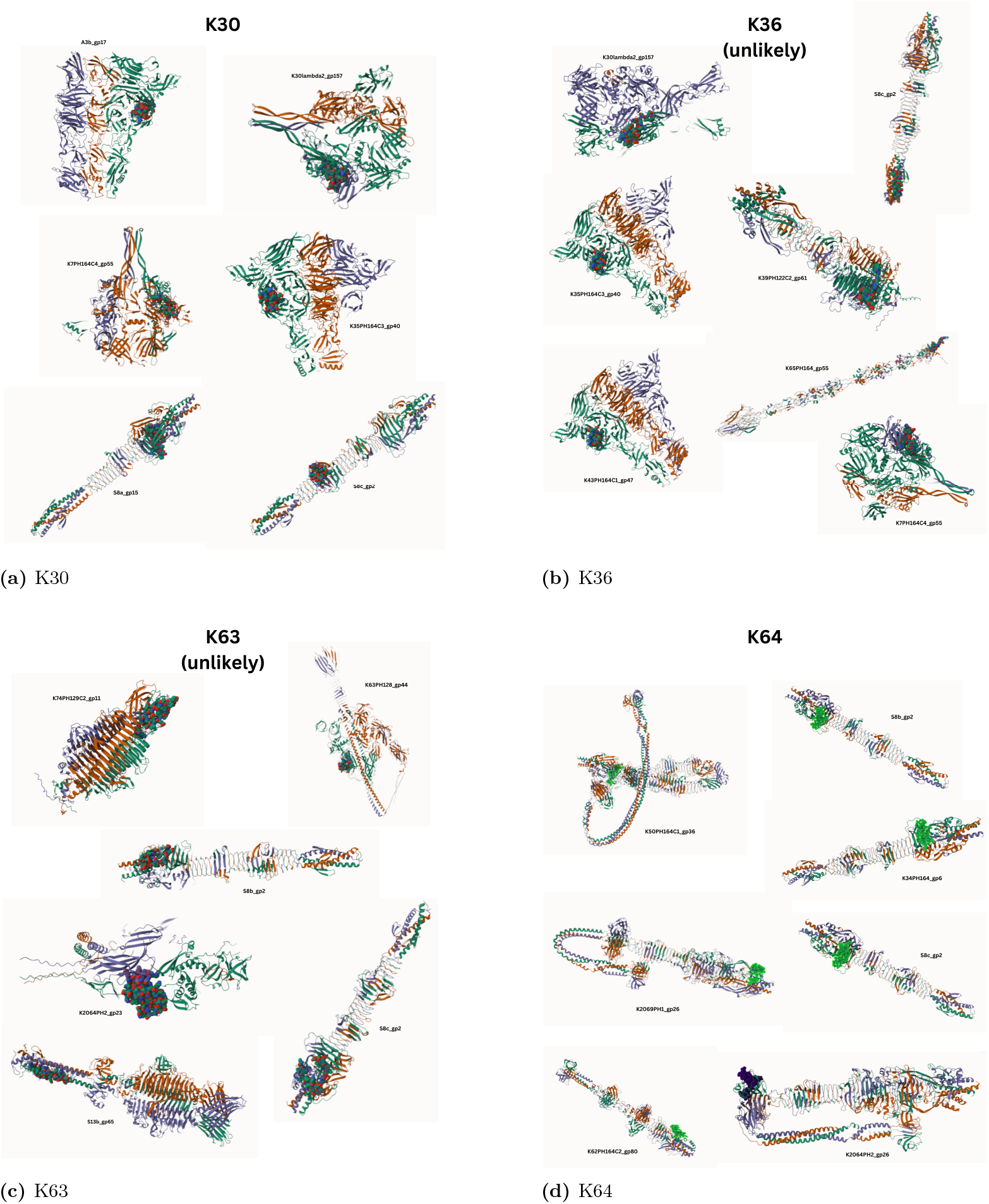
Predicted structures of RBPs containing conserved serotype-associated motifs for K30, K36, K63, and K64. Motifs are shown in molecular representation, replacing the ribbon representation used for the rest of each protein.

**S4 Appendix. Comparison of original and motif-informed protein-level models**. This appendix provides the full comparison between the original protein-level model trained with unmodified interaction labels and the motif-informed relabelled model. These tables support the conclusion that motif-informed relabelling increases score separation among RBPs from the same phage and improves prioritization of motif-bearing candidate adsorption proteins.

**Table 4.** Standard deviation of predicted interaction scores across RBPs encoded by the same phage and targeting the same serotype, grouped by the number of RBPs. The revised model used motif-bearing RBPs as positive protein-level samples.

| Number of RBPs | 2 | 3 | 4 | 5 | 6 | 7 | 8 |
| --- | --- | --- | --- | --- | --- | --- | --- |
| Original model, unmodified labels | .023 | .025 | .078 | .025 | .034 | .009 | .028 |
| Relabelled model, motif-informed labels | .191 | .251 | .196 | .462 | .370 | .261 | .166 |

**Table 5.** Motif-bearing RBPs identified as most informative by the original protein-level model with unmodified interaction labels at the 100% grouping threshold.

| Number of RBPs | 2 | 3 | 4 | 5 | 6 | 7 | 8 |
| --- | --- | --- | --- | --- | --- | --- | --- |
| Number of samples | 37 | 28 | 3 | 42 | 5 | 0 | 1 |
| Motif-bearing RBPs identified as most informative | 15<br>(40.5%) | 10<br>(35.7%) | 3<br>(100%) | 9<br>(21.4%) | 1<br>(20.0%) | X | 0<br>(0.0%) |
| Expectation under random selection | 18.5<br>(50.0%) | 9.33<br>(33.3%) | 0.75<br>(25.0%) | 8.4<br>(20.0%) | 0.83<br>(16.6%) | X | 0.125<br>(12.5%) |
| Binomial test $p$ | .9061 | .4645 | **.0156** | .4691 | .5981 | X | 1.0000 |

**Table 6.**
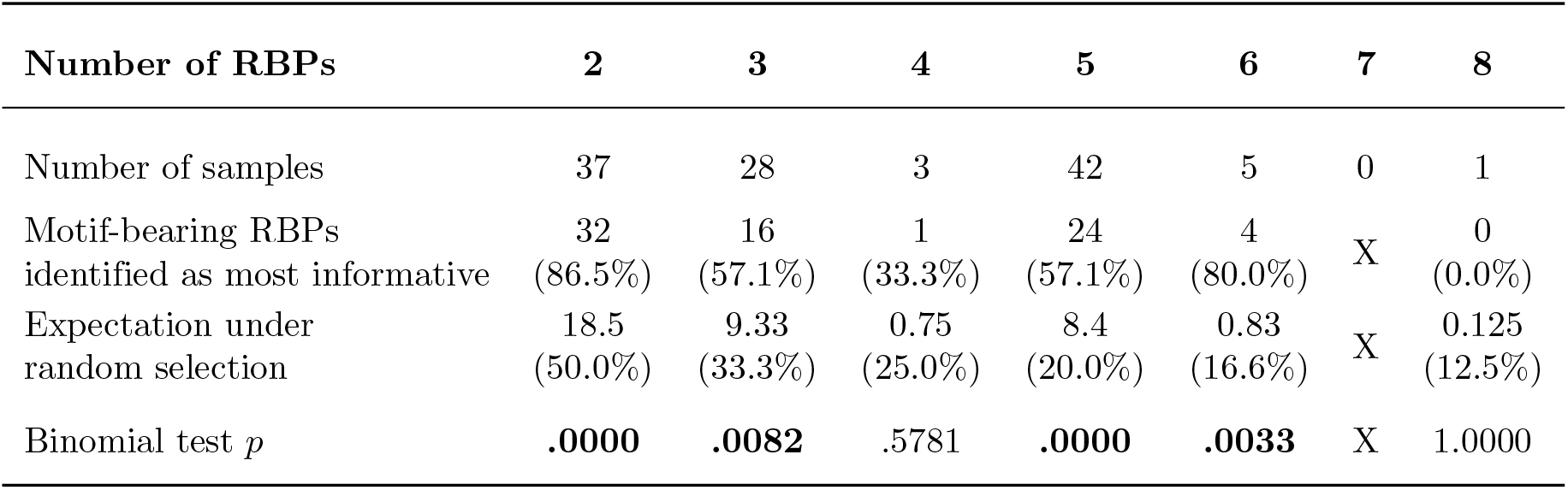
Motif-bearing RBPs identified as most informative by the motif-informed relabelled model at the 100% grouping threshold. Bolded *p*-values indicate statistical significance at the .05 level. X marks an absent value.

| Number of RBPs | 2 | 3 | 4 | 5 | 6 | 7 | 8 |
| --- | --- | --- | --- | --- | --- | --- | --- |
| Number of samples | 37 | 28 | 3 | 42 | 5 | 0 | 1 |
| Motif-bearing RBPs identified as most informative | 32<br>(86.5%) | 16<br>(57.1%) | 1<br>(33.3%) | 24<br>(57.1%) | 4<br>(80.0%) | X | 0<br>(0.0%) |
| Expectation under random selection | 18.5<br>(50.0%) | 9.33<br>(33.3%) | 0.75<br>(25.0%) | 8.4<br>(20.0%) | 0.83<br>(16.6%) | X | 0.125<br>(12.5%) |
| Binomial test $p$ | **.0000** | **.0082** | .5781 | **.0000** | **.0033** | X | 1.0000 |

**S5 Appendix. Full amino-acid and nucleotide sequence of gp2 from bacteriophage S8c**. This appendix reports the amino-acid and nucleotide sequence of gp2 from bacteriophage S8c, the RBP identified as containing motifs associated with multiple capsular serotype groups. Providing the full sequence facilitates independent inspection, replication, and future experimental validation of this candidate multi-motif protein. Individual extracted protein information is also available in the Zenodo repository in the file `RBPbase.csv`.

### Amino-acid sequence

~~~
>S8c_gp2 | receptor-binding protein | length = 414 aa
MANISDQLAADIHNAFNKYYTDLTNQDQIFF
GVGNVTITKQDGTTATVRSWNKVIGSVDTAA
QRGAENNFTSVQTFSAGMKINGNVNCMADNA
MIYLGKNSDLALLKKSGQGGTIAVGSGTPFK
IQRTNTATVSPASTVEDILTIGTDKKTTVAG
ALAAGGDVTAKGNIDNTASGKMYSQALELSF
GTPYIDFHFNNSTADYTTRLIELVSGELTLE
GAFMCKRHLYAWGSLMARSVAPSNPPTGQLI
TGAPFQSMIQGRGANGDARGAVANYYIEELV
GTEHRIVAYLDGYGRSDAWIFRAGGTLTTPK
GDVLTTGSDVRLKTDFTQAPENASERIERLG
VCEYRMKGETRVRRGFIAQQAETVDKVYTYQ
DVEQEIDGERIKVMNVDYVAIIADLVSSVQE
LKAQIRELKGE
~~~

### Nucleotide sequence

~~~
>S8c_gp2 | gene gp2 | length = 1242 bp (excluding stop codon)
ATGGCAAATATTAGCGATCAGCTCGCGGCTG
ATATTCACAACGCGTTTAACAAATACTACAC
AGACCTGACGAACCAAGATCAGATTTTCTTC
GGCGTCGGTAATGTGACAATCACGAAGCAGG
ACGGCACCACCGCCACCGTGCGCTCATGGAA
CAAGGTGATCGGCTCAGTGGACACCGCAGCG
CAGCGCGGGGCGGAGAATAACTTTACCTCAG
TACAGACGTTCAGCGCCGGGATGAAGATTAA
CGGTAACGTCAACTGTATGGCCGACAACGCC
ATGATCTACCTTGGTAAGAACTCAGATCTTG
CGCTGCTTAAAAAGAGCGGGCAGGGTGGAAC
CATCGCCGTCGGCAGCGGCACGCCGTTTAAG
ATCCAGCGCACGAACACCGCCACCGTATCAC
CGGCGTCGACAGTTGAGGATATTCTGACCAT
CGGCACGGATAAGAAAACCACGGTTGCCGGA
GCGCTTGCCGCTGGTGGTGATGTGACGGCGA
AGGGGAACATTGACAACACGGCCAGCGGTAA
GATGTATTCCCAGGCGCTGGAGCTATCATTT
GGCACGCCATACATTGATTTTCATTTCAATA
ACAGCACTGCGGATTACACCACACGGCTGAT
CGAACTGGTATCTGGTGAGCTGACCCTTGAA
GGTGCATTTATGTGCAAAAGGCATCTTTACG
CATGGGGATCGCTTATGGCTCGGTCGGTGGC
ACCAAGCAACCCACCAACCGGGCAACTCATT
ACCGGCGCACCGTTCCAGTCGATGATCCAGG
GGCGAGGAGCAAACGGTGATGCGCGCGGCGC
GGTGGCTAACTACTACATCGAGGAACTGGTT
GGAACCGAGCACCGGATCGTCGCTTACCTCG
ACGGATACGGGAGATCCGACGCGTGGATCTT
CCGGGCTGGTGGCACGCTCACCACGCCGAAG
GGTGACGTCCTAACGACCGGTTCCGACGTGC
GCCTGAAAACAGACTTCACGCAAGCGCCTGA
AAACGCCTCAGAGCGCATTGAGCGCTTAGGG
GTATGTGAGTACCGGATGAAGGGAGAAACGC
GCGTTAGGCGTGGTTTTATCGCGCAGCAGGC
GGAGACCGTAGACAAGGTCTATACCTATCAG
GACGTCGAGCAGGAAATTGACGGCGAGCGTA
TCAAGGTGATGAACGTCGACTATGTGGCGAT
CATTGCTGACCTTGTGTCATCGGTGCAGGAA
TTGAAGGCGCAGATCAGGGAATTGAAAGGGG
AATGA
~~~

